# Specific ATPases drive compartmentalized glycogen utilization in rat skeletal muscle

**DOI:** 10.1101/2021.10.27.466064

**Authors:** Joachim Nielsen, Peter Dubillot, Marie-Louise H. Stausholm, Niels Ørtenblad

## Abstract

Glycogen is a key energy substrate in excitable tissue and especially in skeletal muscle fibers it contributes with a substantial, but also local energy production. A heterogenic subcellular distribution of three distinct glycogen pools in skeletal muscle is proved by transmission electron microscopy (TEM), which is thought to represent the requirements for local energy stores at the subcellular level. Here, we show that the three main energy-consuming ATPases in skeletal muscles (Ca^2+^-, Na^+^,K^+^-, and myosin ATPases) utilize different local pools of glycogen. These results clearly demonstrate compartmentalized glycogen metabolism and emphasize that spatially distinct pools of glycogen particles act as energy substrate for separated energy requiring processes, suggesting a new paradigm for understanding glycogen metabolism in working muscles, muscle fatigue and metabolic disorders.

## Introduction

It is of vital importance for all cell types to balance energy utilization and production, but it is particularly important in excitable cells with high fluctuations in energy turnover. A close match between energy utilization and production is established by functional compartmentalization of enzymatic reactions^1,2^, which ensures rapid exchange of metabolites. In skeletal muscle, this has probably evolved as a result of both a spatial distribution of energy consuming and producing processes and a physical constraint imposed by intracellular structures limiting free diffusion^3^. Intriguingly, glycogen particles and glycogenolytic and glycolytic enzymes are observed adjacent to the sarcoplasmic reticulum (SR) membrane^4,5^ and it is well described how this physical association directs a crosstalk, where Ca^2+^ release from the SR upon muscle activation facilitates glycogen degradation and glycolytic ATP production and, reversely, where glycogen loss impairs SR function^6,7^. The glycogen-glycogenolytic-glycolytic system is an example of functional compartmentalization, where the specific organization and localization of enzymes and glycogen particles create an efficient delivery of energy to energy requiring processes located in specific subcellular domains^8,9^. Thus, the diffusional barrier by intracellular structures is circumvented by situating the metabolic machinery and the glycogen particles close to the energy consuming processes, where one pool of glycogen particles may only serve the neighboring ATPases.

In working skeletal muscle fibers, myosin ATPases, SR Ca^2+^ ATPases and Na^+^,K^+^-ATPases consume approximately 50-60%, 40-50% and 5-10% of the energy, respectively^10,11^. In order to clarify whether these three major energy consuming processes utilize different pools of glycogen particles, we conducted two sets of experiments, where we stimulated or inhibited specific energy consumption by the different ATPases combined with measures of the distinct pools of glycogen particles by quantitative TEM.

## Results and discussion

We defined three pools of glycogen based on their spatial distribution (Fig. 1A-C). To embrace fiber-to-fiber variation, we estimated the volumetric content at the single fiber level, where coefficient of errors between 0.15 and 0.25 could be obtained after analyses of 12-16 images per fiber (Fig. 1D). The TEM estimated total glycogen volume fraction correlated well with biochemically determined mixed glycogen concentration from homogenates (Fig. 1E). In the resting control muscles combined from both experiments, glycogen particles were distributed with 65%, 29% and 6% as intermyofibrillar, intramyofibrillar and subsarcolemmal glycogen, respectively (Fig. 1F). This pattern is consistent with a previous study in rats (*12*), but the share of intramyofibrillar glycogen in rat and mouse skeletal muscle is ∼3-fold larger than in humans, due to a lower absolute amount of intermyofibrillar glycogen in rats and mice^13,14^.

**Fig. 1.**
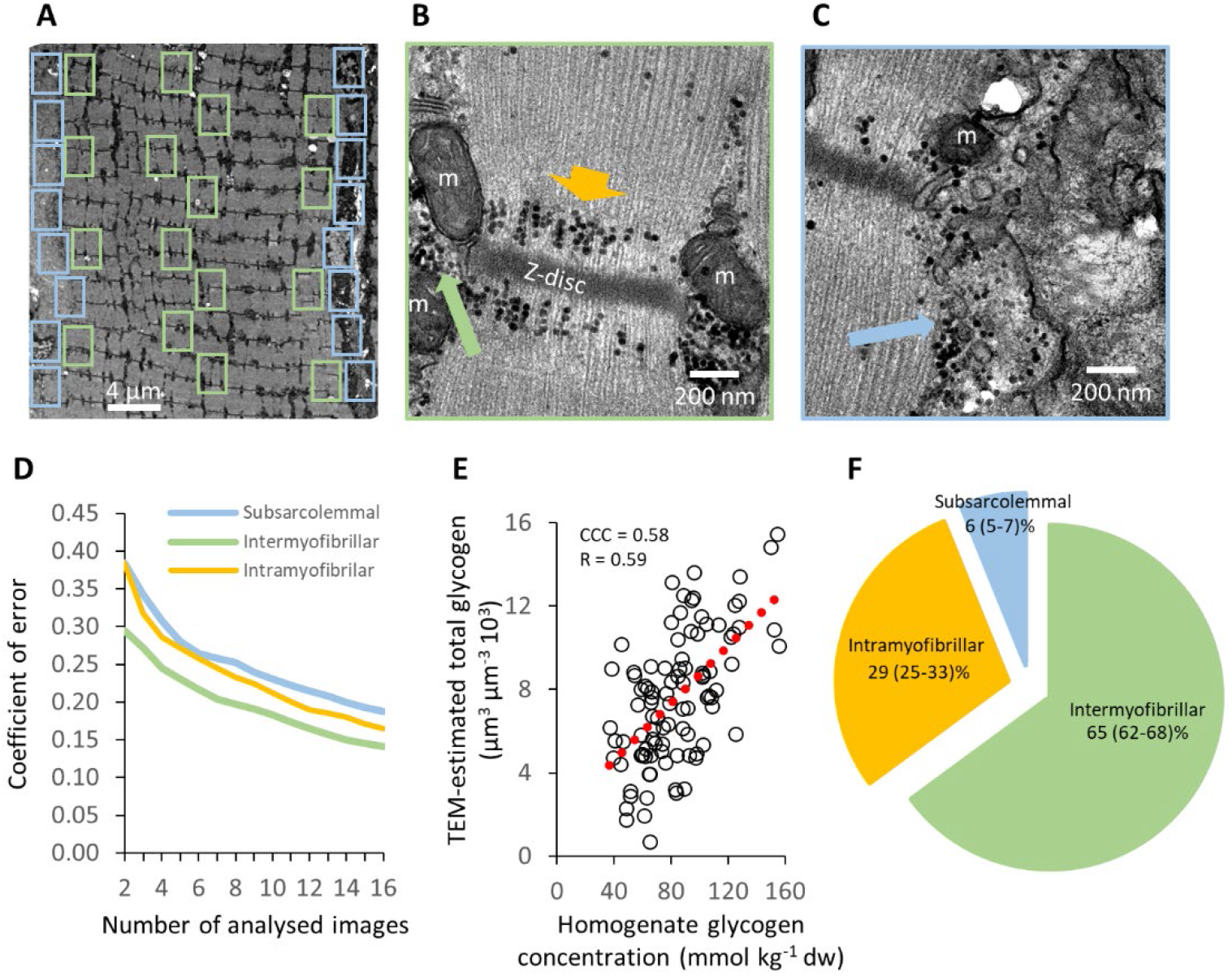
TEM analyses of subcellular glycogen distribution. **(A)** Fixed muscle segments were prepared for glycogen visualization by TEM and cut in longitudinal sections. Three fibers were photographed per muscle in a randomized systematic order including 12-16 images from the subsarcolemmal region and 12-16 images from the myofibrillar region. **(B)** In the myofibrillar images, the volumetric content of intermyofibrillar glycogen (long green arrow) and intramyofibrillar glycogen (short orange arrow) were estimated by point counting (*13, 26*). **(C)** In the subsarcolemmal images, the volume of subsarcolemmal glycogen (blue arrow) per fiber surface area was estimated by point counting and a fiber length measurement (*13*). **(D)** After completion of analysis of the first 102 fibers, image-to-image variation showed a mean stereological coefficient of error of 0.14, 0.17 and 0.19 after analysis of 16 images for intermyofibrillar, intramyofibrillar and subsarcolemmal glycogen, respectively. It was decided for the remaining fibers (n= 172) that 12 photographed images were sufficient per region. **(E)** Scatterplot of TEM-estimated total glycogen per muscle (mean of three fibers) versus the glycogen concentration determined from a homogenate. Condordance correlation coefficient (CCC) showed a moderate agreement between the two methods. R indicates Person’s correlation coefficient. Dotted red line indicates best linear fit. (**F**) The relative distribution (%) of the three pools to the total volumetric content. Values are mean and 95% confidence interval. *n* = 60 fibers from 20 rats.

In the first experiment, we found that by blocking the myosin ATPase by N-benzyl-p-toluene sulfonamide (BTS)^15^ and blebbistatin^16^ without affecting excitability^17^, the tetanic stimulation-induced force production is minimal (Fig. 2A) accompanied by less utilization of mixed muscle glycogen (Fig. 2B) and less production of lactate (Fig. 2C). By blocking the myosin ATPase, the utilization of intramyofibrillar glycogen was almost completely abolished, while the utilization of intermyofibrillar glycogen was reduced by about 50% (Fig. 2D-E). Since the intermyofibrillar pool contains twice the amount of glycogen than the intramyofibrillar pool, the myosin ATPase taxes those two pools equally in absolute quantity. When the myosin ATPase is blocked, the SR Ca^2+^ ATPase represent the majority (>80%) of the remaining energy consumption during contractions, of which the requirement for glycogen therefore can be exclusively connected to the utilization of intermyofibrillar glycogen. This differential glycogen utilization by the myosin ATPases and SR Ca^2+^ ATPases is in line with the idea of a functional compartmentalization of energy production and utilization in the muscle cell and the proximity of the pools and the specific ATPases (Fig. 2F-G).

**Fig. 2.**
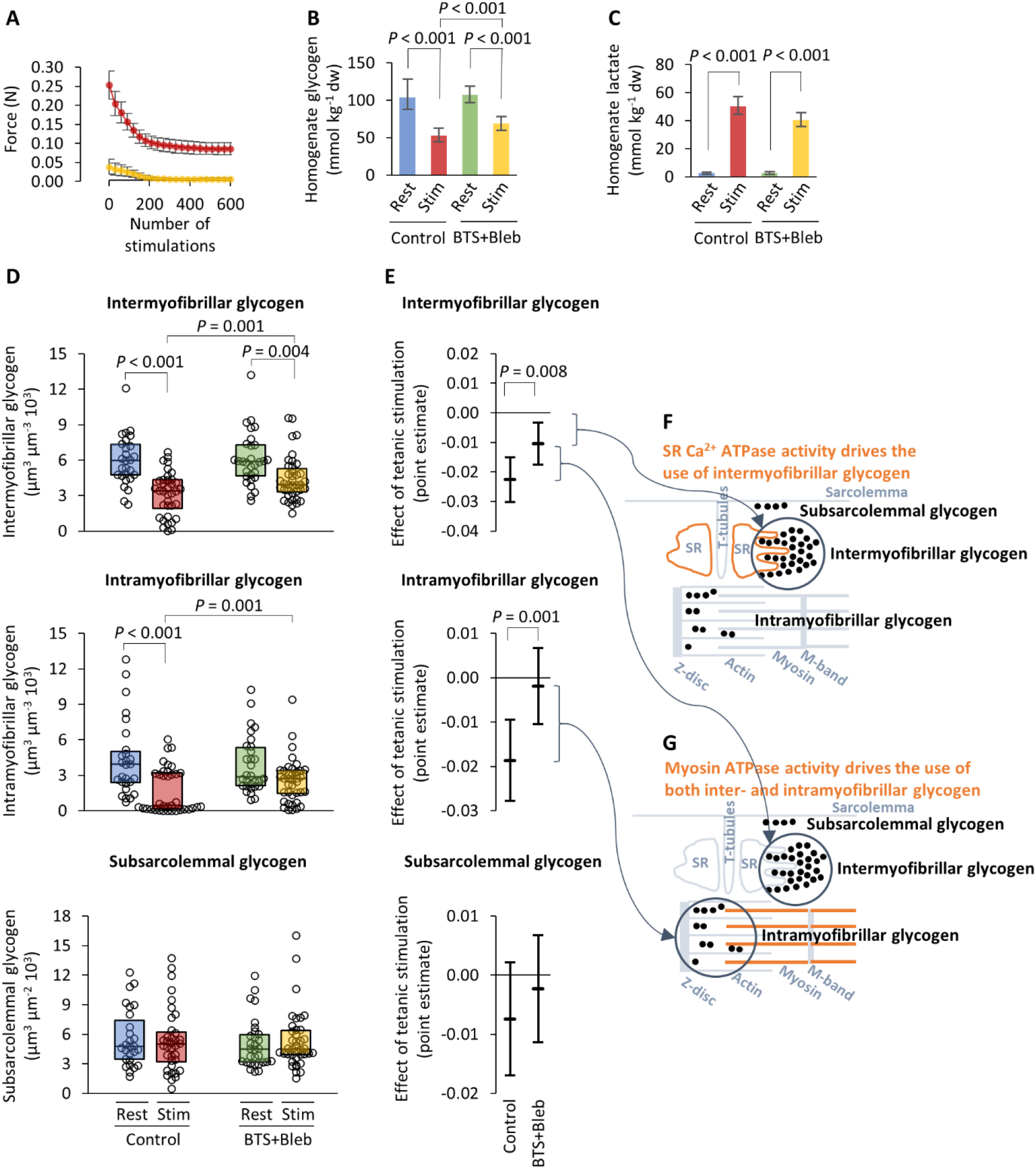
Effect of myosin ATPase inhibition during tetanic stimulation on subcellular compartmentalized glycogen metabolism. Soleus muscles of Sprague Dawley rats (4-6-week-old males weighing 112-276 g, killed by cervical dislocation) were mounted to a force transducer and bathed in a standard Krebs-Ringer bicarbonate buffer with 4 mM K^+^ and 5 mM D-glucose (pH 7.4) at 30°C continuously gassed with a mixture of 95% O_2_ and 5% CO_2_. After 30 min of rest in the standard KR buffer, the muscles were incubated for 180 min with either 50 µM N-benzyl-p-toluene sulfonamide (BTS) and 25 µM blebbistatin or vehicle (DMSO; 0.3 volume%). Then, muscles were either tetanic stimulated (30 Hz for 400 ms every 2 sec) or rested for 20 min. (**A**) Force production during repeated tetanic stimulations in control muscles (red) and in muscles with the myosin ATPase inhibited (BTS+Bleb) (yellow) shown as mean and SD. *n* = 12-13 muscles. (**B**) Muscle homogenate glycogen concentration shown as mean with 95% confidence interval. *n* = 9-13 rats. (**C**) Muscle homogenate lactate concentration shown as geometric mean with 95% confidence interval. *n* = 9-13 rats. Individual values are presented in Figure 1S. (**D**) Single fibre values of glycogen in three distinct subcellular pools: intermyofibrillar, intramyofibrillar and subsarcolemmal. Data are shown as box plots displaying the first and third quartiles and split by the median. *n* = 27-39 fibers from 9-13 muscles. (**E**) Point estimates with 95% confidence interval from linear mixed effect model on square root-transformed data from (D). (**F-G**) Illustrations of the spatial association between SR Ca^2+^ ATPases situated at the SR membrane (orange in F) and intermyofibrillar glycogen, and between the myosin ATPases (orange in G) and inter and intramyofibrillar glycogen. Interaction or main effects were tested using linear mixed effect model on square root-transformed data with tetanic stimulation and myosin ATPase inhibitors as fixed effects and rat ID as random effect. *P* values for two-way interactions were 0.008, 0.001, 0.395, 0.039 and 0.045 for intermyofibrillar, intramyofibrillar, subsarcolemmal glycogen, homogenate glycogen and homogenate lactate, respectively.

As the Na^+^,K^+^-ATPase only consumes 5-10% of the total energy consumption during work^11^, we needed to take another approach to investigate it’s potential connection to a local pool of glycogen. In this second experiment, muscles were exposed to the β-adrenoceptor agonist salbutamol, which stimulate the Na^+^,K^+^-ATPase^18^, combined with the cardiac glycoside ouabain, which selectively block the Na^+^,K^+^-ATPase^19^. Here, salbutamol mediates a reduction in mixed glycogen concentration and an increase in lactate concentration, which can be attenuated by ouabain (Fig. 3D-E) corroborating a previous study on mixed glycogen concentration^20^.

**Fig. 3.**
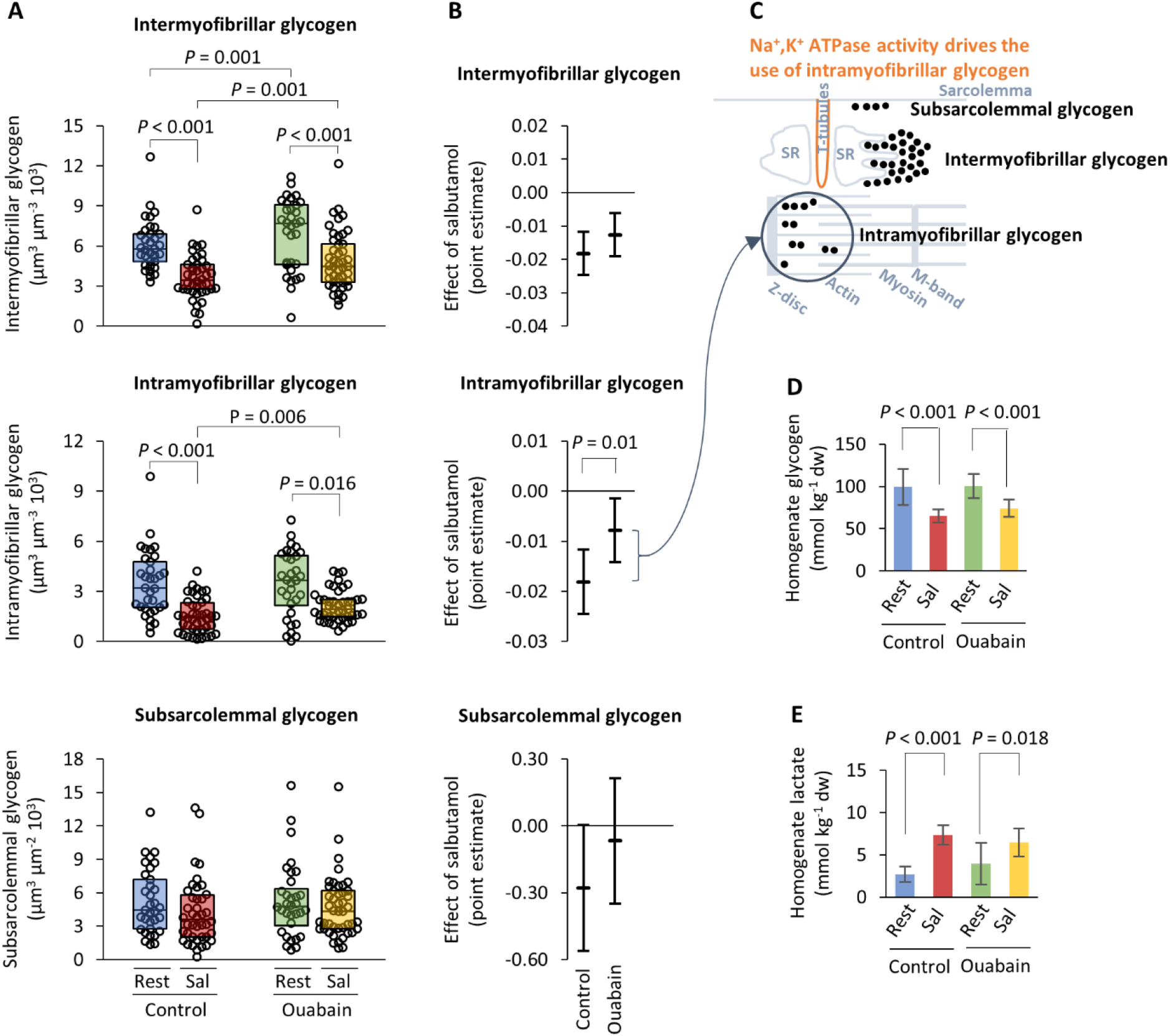
Effect of Na^+^,K^+^ ATPase inhibition during salbutamol exposure on subcellular compartmentalized glycogen metabolism. Soleus muscles of Sprague Dawley rats (4-6-week-old males weighing 102-192 g, killed by cervical dislocation) were mounted to a force transducer and bathed in a standard Krebs-Ringer bicarbonate buffer with 4 mM K^+^ and 5 mM D-glucose (pH 7.4) and 1 mM ouabain or vehicle (water) at 30°C continuously gassed with a mixture of 95% O_2_ and 5% CO_2_. After 30 min of rest, the muscles were incubated for 60 min with either 10 µM salbutamol or vehicle (methanol; 2 volume%). (**A**) Single fiber values of glycogen in three distinct subcellular pools: intermyofibrillar, intramyofibrillar and subsarcolemmal. Data are shown as box plots displaying the first and third quartiles and split by the median. *n* = 32-45 fibers from 11-16 muscles. (**B**) Point estimates with 95% confidence interval from linear mixed effect model on square root-transformed data from (intermyofibrillar and intramyofibrillar glycogen) or log-transformed data (subsarcolemmal glycogen) from (A). (**C**) Illustrations of the spatial association between Na^+^,K^+^ ATPases situated at the t-tubular membrane (orange) and intramyofibrillar glycogen. (**D**) Muscle homogenate glycogen concentration shown as mean with 95% confidence interval. *n* = 11-16 muscles. (**E**) Muscle homogenate lactate concentration shown as mean with 95% confidence interval. *n* = 11-16 muscles. Individual values are presented in Figure 2S. Interaction or main effects were tested using linear mixed effect model on square root-transformed (intermyofibrillar and intramyofibrillar glycogen in A), log-transformed (subsarcolemmal glycogen in A) or non-transformed data (D and E) with salbutamol and ouabain as fixed effects and rat as random effect. *P* values for two-way interactions were 0.164, 0.010, 0.225, 0.281 and 0.052 for intermyofibrillar, intramyofibrillar, subsarcolemmal glycogen, homogenate glycogen and homogenate lactate, respectively.

However, ouabain treatment induced a large reduction in the salbutamol-induced utilization of intramyofibrillar glycogen, but no or to a smaller extent reductions in intermyofibrillar and subsarcolemmal glycogen (Fig. 3A-B), demonstrating that the activity of Na^+^,K^+^-ATPase is connected to the utilization of intramyofibrillar glycogen. Since Na^+^,K^+^-ATPase activity occurs in the t-tubular system and the sarcolemma, it is not co-localized with intramyofibrillar glycogen (Fig. 3C). However, glycolytic enzymes are abundant in the t-tubular membrane^21^ creating the structural basis for a functional compartmentalization with distant glycogen particles. A possible connection between intramyofibrillar glycogen and processes occurring in the triadic gap between the SR and the t-tubular system is supported by our previous findings showing an association between intramyofibrillar glycogen and both excitability^12^ and SR Ca^2+^ release rate^7^.

While channeling of metabolites occurs both from glycolysis and mitochondrial oxidative phosphorylation (via the creatine and adenylate kinases phosphotransfer) to all sites of the main ATPases and is more efficient than free diffusion^2^, the glycolytic derived ATP is the key regulator of several energy requiring og sensing processes within skeletal^20^, smooth^8^ and heart muscle^22^, and in the brain^23^. We suggest that a competition between the myosin ATPases and the Na^+^,K^+^-ATPases for intramyofibrillar glycogen creates a link, where contractility is connected to excitability. This explains why muscles devoid of glycogen due to a prior sustained high consumption rate or inherited diseases suffer from muscle fatigue^24^ and exercise intolerance^25^.

## Methods

### Animals

All handling and use of animals complied with Danish animal welfare regulations.Experiments were performed using 4-6 week-old Wistar rats of own breed, weighing 102-276 g, which were kept in a thermostated environment at 21 °C with a 12 h / 12 h light - dark cycle and fed ad libitum at the Biomedical Laboratory, University of Southern Denmark.

### Force measurements

Muscles were mounted for isometric contractions in thermostated chambers containing standard KR buffer and adjusted to optimal length for force production. Force was measured using force displacement transducers and recorded with a chart recorder and digitally on a computer.

### Krebs Ringer solution for In vitro incubation of soleus muscles

All in vitro experiments were performed with muscles incubated in a standard Krebs Ringer bicarbonate buffer containing the following (in mM) 122.1 NaCl, 25.1 NaCHO_3_, 2.8 KCl, 1.2 KH_2_PO_4_, 1.2 MgSO_4_, 1.3 CaCl_2_, and 5.0 D-glucose (pH 7.4).

### Chemicals

All chemicals were of analytical grade and unless stated were obtained from Sigma-Aldrich, with BTS obtained from Toronto Research Chemicals, Ontario, Canada.

### Determination of muscle glycogen and lactate concentration

Homogenate glycogen concentration was determined by spectrophotometry (Beckman DU 650). Freeze dried muscle tissue (1.5 mg) was boiled in 0.5 ml 1 M HCL for 150 min. before it was rapidly cooled, whirl-mixed and centrifuged at 3500g for 10 min. at 4° C. 40 μL of boiled muscle sample and 1 ml of reagent solution containing Tris-buffer (1M), distilled water, ATP (100mM), MgCl2 (1M), NADP^+^(100mM) and G-6-PDH were mixed before the process was initiated by adding 10μL of diluted hexokinase. Absorbance was recorded for 60 min. before the glycogen content was calculated. Lactate was determined from specimen, which was freeze-dried, dissected free of non-muscle tissue, powdered and extracted with HClO4. Muscle glycogen and lactate was expressed as mmol·kg^-1^ dw.

### Quantitative transmission electron microscopy

#### Preparation of samples for glycogen staining

A small piece (< 1mm^3^) of the mid belly of m. soleus was fixed in 2.5 % glutaraldehyde in 0.1 M sodium cacodylate buffer (pH 7.3) for 24 h at 4° C and subsequently rinsed four times in 0.1 M sodium cacodylate buffer. Following rinsing, the muscle pieces were post-fixed with 1 % osmium tetroxide (OsO4) and 1.5 % potassium ferrocyanide (K4Fe(CN)6) in 0.1 M sodium cacodylate buffer for 90 min at 4° C. The use of potassium ferrocyanide during postf fixation enhance the visualization of glycogen particles. After post-fixation, the muscle pieces were rinsed twice in 0.1 M sodium cacodylate buffer for 60 min at 4° C, dehydrated through a graded series of alcohol at 4–20° C, infiltrated with graded mixtures of propylene oxide and Epon at 20° C, and embedded in 100 % Epon at 30° C. Ultra-thin (60 nm) sections were cut (using a Leica Ultracut UCT ultramicrotome) in two depths (separated by 150 μm) and contrasted with uranyl acetate and lead citrate. Sections were examined and three longitudinal oriented fibers per muscle were photographed in a pre-calibrated transmission electron microscope (JEM-1400Plus, JEOL Ltd, Tokyo, Japan and a Quemesa camera or Philips CM100 and an Olympus Veleta camera). Images were analyzed using a digital screen at a final total magnification of x100,000.

#### Quantification of the subcellular distribution of glycogen

The volume fraction of glycogen in three distinct localizations was estimated using standard stereological techniques. Since glycogen particles (diameter of 10-40 nm) are smaller than the thickness of the section (60 nm), the calculation of the volume fraction based on an area fraction on the projected images is corrected for an overestimation due to a cutting by the upper and lower slice surface and thereby abrogating the spherical shape of some of the particles. This is done by the formula suggested by Weibel^26^:

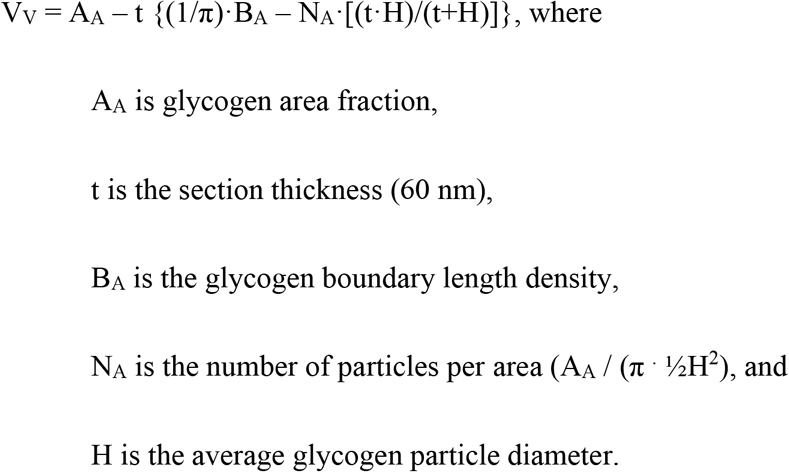

Glycogen particles were assumed to be spherical. A_A_ was estimated by point counting using different grid sizes for the different locations in order to achieve satisfactorily precision of the estimates (see below).

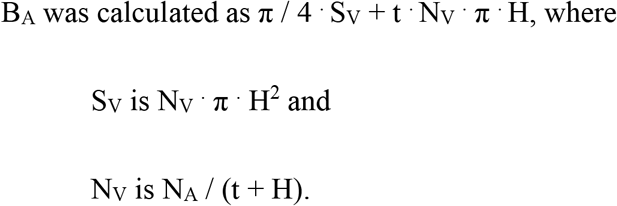

The average glycogen particle diameter for each location was calculated by directly measuring at least 60 particles per location per fiber using iTEM (iTEM software, version 5.0, Olympus, Germany). However, of the 284 fibers analyzed, only 40 to 59 particles could be found in 72 fibers and only 10 to 39 particles in 24 fibers. Intermyofibrillar glycogen was expressed relative to the myofibrillar space and estimated using grid sizes of 120 nm and 300 nm, respectively. The amount of intramyofibrillar glycogen was expressed relative to the intramyofibrillar space and estimated using grid sizes of 60 nm and 300 nm, respectively. The subsarcolemmal glycogen was expressed relative to the muscle fiber surface area and estimated using a grid size of 90 nm. The fiber surface area was estimated by measuring directly the length of the fiber accompanying with the area of the subsarcolemmal region, which is perpendicular to the outer most myofibril and then multiplied by the section thickness (60 nm).

## Acknowledgments

The experiments were performed at the Institute of Sports Science and Clinical Biomechanics (in vitro experiments and metabolite analyses) and Institute of Pathology, Faculty of Health Science (transmission electron microscopy analyses), University of Southern Denmark, DK-5230 M, Denmark. We thank Kirsten Hansen and Karin Trampedach for skillful technical assistance. This study was supported by a grant from The Ministry of Culture Committee on Sports Research (TKIF2011-058).

## Author contributions

J.N. contributed with conceptualization, formal analysis, investigation, data curation, writing – original draft, visualization, supervision, project administration, and funding acquisition. P.D. contributed with conceptualization, formal analysis, investigation, writing – review & editing, visualization. M.L.H.S. contributed with investigation, writing-review & editing. N.Ø. contributed with conceptualization, investigation, writing – review & editing, supervision.

